# Rapid evolution of the functionally conserved gap gene *giant* in *Drosophila*

**DOI:** 10.1101/2021.07.08.451553

**Authors:** Wenhan Chang, Daniel R. Matute, Martin Kreitman

## Abstract

Developmental processes in multicellular organisms, and the outcomes they produce, are often evolutionarily conserved. Yet phylogenetic conservation of developmental outcomes is not reflected in functional preservation of the genes regulating these processes, a phenomenon referred to as developmental system drift (1, 2). Little is known about the evolutionary forces producing change in the molecular details of regulatory genes and their networks while preserving development outcomes. Here we address this void in knowledge by systematically swapping the *Drosophila melanogaster* coding and noncoding regions of the essential gap gene, *giant*, a key regulator of embryonic pattern formation, with orthologous sequences drawn from both closely and distantly related species within the genus. Employing sensitized genetic complementation assays, the loss of a transgene’s ability to restore viability occurs across phylogeny at every interspecific level of comparison and includes both coding and noncoding changes. Epistasis is present as well — both between coding and noncoding sequences and, in a dramatic example of change-of-sign epistasis, between the only two coding substitutions separating two very closely related species. A continuous process of functional divergence hidden under conserved phylotypic developmental outcomes requires reconsideration of the prevailing view that the essential genes in conserved regulatory networks are protected from the driving forces of evolutionary change.

## Introduction

The preservation of molecular function is a universal theme in the evolution of life, evident in the myriad of recognizably conserved molecules, proteins, genetic pathways and biochemical processes across phylogeny. All multicellular organisms, for example, possess a shared set of Hox genes regulating cell differentiation and development (3). Conserved molecular and gene expression phenotypes are believed to reflect intricately buffered developmental pathways that constrain functional evolution of member genes and circuits (4, 5). Support for this view is dominated by experiments emphasizing partial activity or replaceability of a *Drosophila* gene with transgenes carrying orthologs from species as distant as chicken or even human (6–9). Yet, these orthologs (13 instances in total) never fully rescue the mutant phenotypic, and they also do not restore viability (Table S1). Conservation of developmental outputs might belie functional changes in molecules that govern those outputs (10, 11).

Instances of this tension is apparent in the *Drosophila* gap gene network, a set of exquisitely studied transcription factors expressing early in embryonic development to orchestrate the highly conserved process of insect pattern formation (12). Spatio-temporal expression of the gap genes are remarkably conserved across *Drosophila* phylogeny, measured at nuclear resolution in three dimensions and time (13). So too is the cis-regulatory output of the pair-rule gene *even-skipped*, a primary target of the gap genes. When placed in *D. melanogaster, eve* enhancers from species in family Sepsidae, a sister group to *Drosophila*, respond to *D. melanogaster* gap proteins by driving pair-rule stripe expression nearly identically to the native *eve* expression pattern (14, 15), this despite extensive rearrangement of the relevant transcription factor binding sites. In contrast, other insect taxa, including mosquitos and moth fly, employ different maternal genes to establish head-to-tail polarity (16). In the scuttle fly, the initiation and expression of the gap genes are, moreover, quantitatively different than *Drosophila*, though the embryos converge to a similar developmental phenotype (17, 18).

These scattershot observations underscore the lack of a mechanistic basis for interpreting developmental system drift and highlight the need for careful systematic measurements of regulatory gene functional divergence across a phylogeny. Do these genes evolve? Is functional divergence compartmentalized to changes in cis-regulation, or do the transcription factors evolve as well? And, if so, what is the evolutionary timescale (and phylogenetic consistency) of change? We focused our experimental investigation on the gap gene *giant* (*gt*) across six *Drosophila* species whose phylogenetic ancestries range from about 1 million years ago (MYA) to about 40 MYA (19, 20). The Giant protein (Gt), a basic leucine zipper transcription factor, is among the earliest proteins expressed zygotically in the blastoderm *Drosophila* embryo to establish landmarks for anterior-posterior patterning and segmentation (21, 22). Its role as a gap gene is conserved over 350 million years of divergence in Oncopeltus (23), and its DNA-binding domain remains extensively conserved in *Drosophila* (Fig. S1) and across bilateria evolution (24). Here we document the pace of *giant* functional divergence in *Drosophila*, both for coding and noncoding regions of the locus, and provide a mechanistic framework for understanding developmental system drift — how a regulatory network can evolve at the molecular level while maintaining a conserved system output.

## Results

### xperimental approach

We employed phiC31 site-specific genetic transformation (25) to study the phenotypic output of *giant* alleles from different species when placed in *D. melanogaster* (*mel*) (Fig. 1). We generated *gt* whole-locus genotypes carrying sequences orthologous to the 27kb native locus — an interval that restores viability in a complementation assay with the *mel* control transgene (26) (Fig. 1b, c). We also generated interspecies transgene chimeras by swapping with the *mel* whole-locus sequence either the protein coding or the noncoding region from each of the five other species (Table S2; a total of 30 transgenic lines). The fluorescent protein eGFP is commonly appended to proteins as a tag to visualize their cellular distribution and function (27). As a means for amplifying possible functional differences among *gt* proteins, we added, in a parallel set of transgenes, an eGFP carboxy-terminal tag to our whole-locus and chimeric transgenes. We scored the relative viability (hereafter RV) — defined as the ratio of F1 flies carrying either an interspecies or control transgene, identified with fluorescent eye markers — in the offspring of test crosses carrying a null allele (*gt*^X11^) at the native locus (Fig. S2, S3). We measured RV in both male and female separately, anticipating that transgene restoration of viability by *gt* orthologs might differ in the two sexes (26). We also sensitized our RV measurements by analyzing flies carrying a single copy of the *gt* transgene.

**Figure 1.**
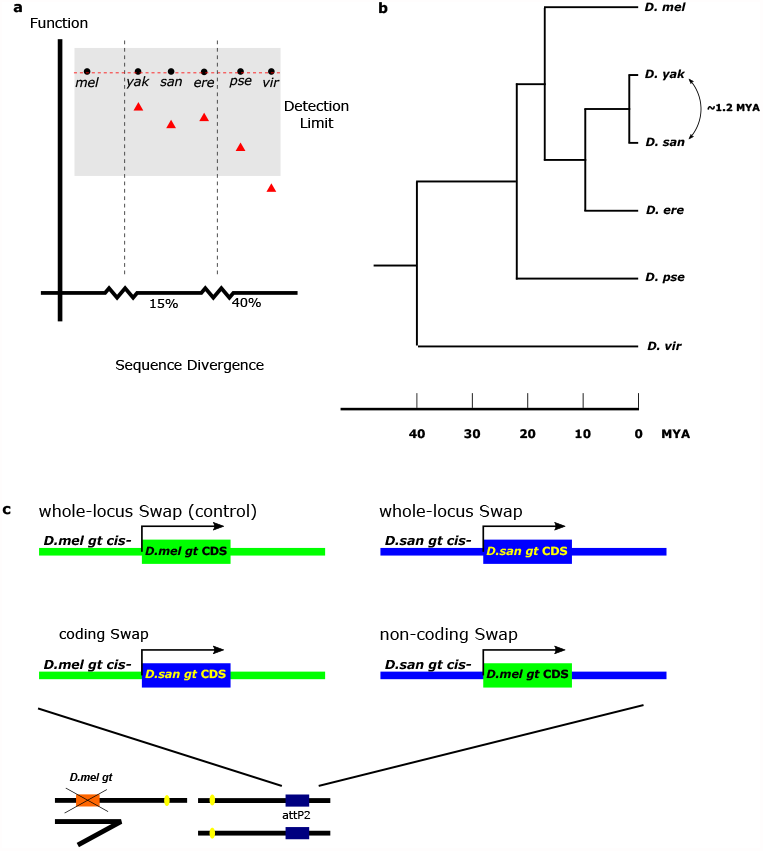
Approach to testing for functional divergence of *gt* transgene orthologs. **a**, Competing hypotheses: Functional stasis (black dots) - *gt* orthologs will be indistinguishable; Functional divergence (red triangles) - *gt* orthologs diverge and will be distinguishable if experimental design has sufficient resolution. In this example, only *vir gt* functional divergence is detectable (shaded region depicts limits of experimental resolution). Vertical dashed lines mark sequence divergence relative to *mel* (Methods). **b**, Phylogenetic relationship of species investigated. **c**, Site-specific phiC31 transgenesis using whole-locus and chimeric *giant*.

### Functional divergence of distant orthologs

We first investigated the *gt* ortholog from *D. virilis* (*vir*), the most distant relative in the genus to *mel* (Fig. 1b; common ancestor 40 MYA (19)). Whole-locus RV is significantly reduced (RV = 0.56) in males and is essentially lethal in females (Fig 2a, d, g). The *vir* coding region alone restores full RV in both sexes; *vir* noncoding sequences restores full RV in males but reduces RV significantly in females, though not to lethality ((RV = 0.21); Fig 2f). The lethality driven by whole-locus *vir* in females, therefore, requires epistatic contributions from the *vir* noncoding and coding regions (Fig 2f). A *vir* coding contribution to loss of RV is confirmed by the eGFP-tagged version of *vir* coding (RV = 0.52; Fig 2h). We also observed a skewed sex ratio in adults when endogenous *gt*^*mel*^ is replaced by two copies of *gt*^*vir*^ (male:female = 3.68:1, Table S7). Collectively, these results identify functional differences in both coding and noncoding regions and reveal epistasis for RV between the two regions.

**Figure 2.**
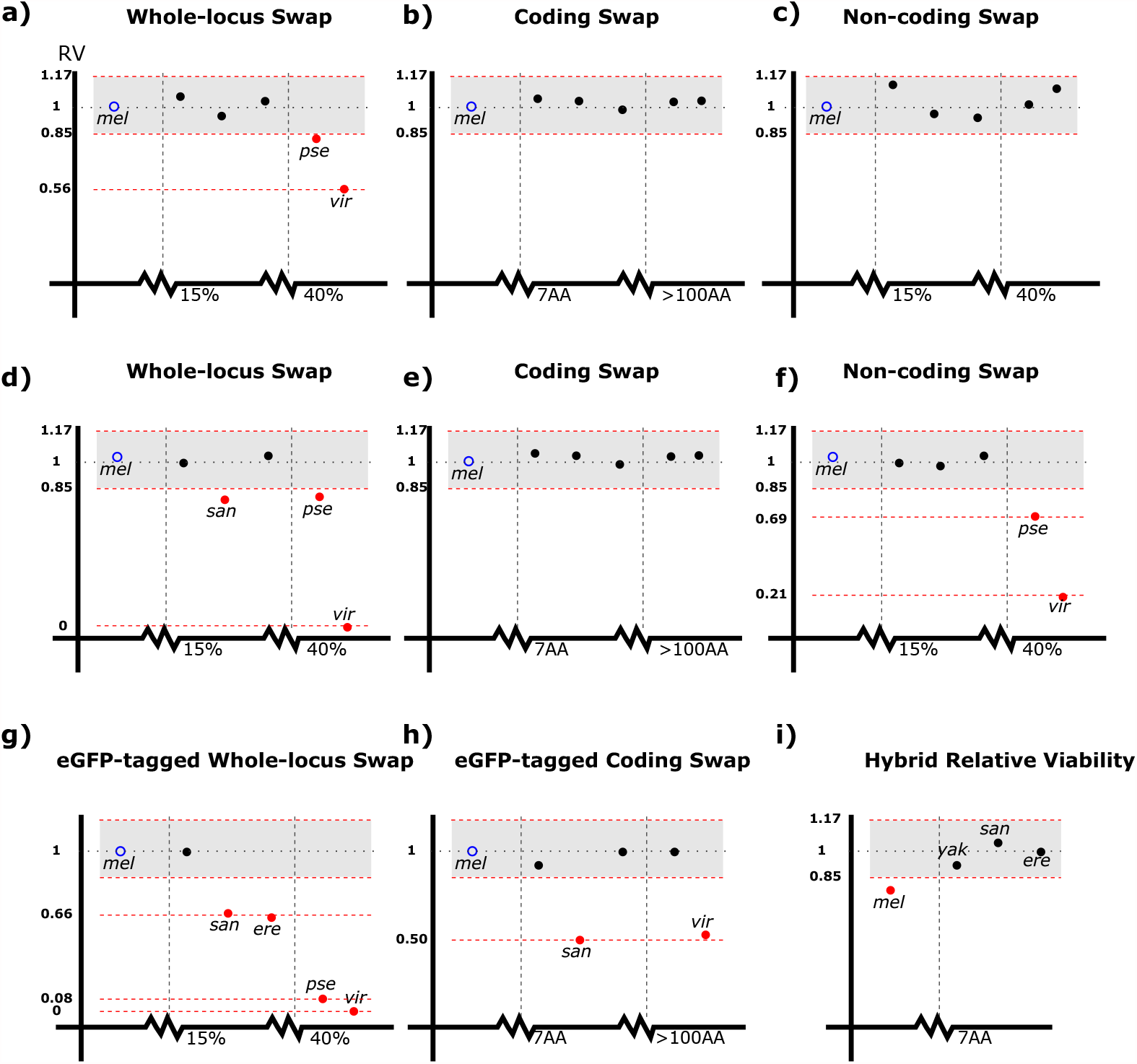
Complementation assays reveal extensive functional divergence across *Drosophila* phylogeny. Single copy transgene rescue in males (**a-c**) and females (**d-h**). **d**, Whole-locus *vir gt* restores *mel* female viability at a low rate (<0.2% RV). **g-h**, Female RV using eGFP-tagged *gt* transgenes. Whole-locus eGFP-tagged *vir gt* restores *mel* female viability only when two copies are present. **i**, *mel gt* coding region is deleterious in *mel/san* hybrids. For **a-h**, the species order is (left-right) *mel, yak, san, ere, pse* and *vir*. RV values significantly different from *mel* are labeled in red. For illustrative purpose, shaded region represents 80% power to detect at p<0.05 a 15% difference in viability between control and experimental transgene (e.g., RV=0.85) for a sample of 1240 adults (see Methods). Sample sizes are given in Table S5.

Next, we investigated the *gt* ortholog of *D. pseudoobscura* (*pse*). This species is estimated to share a common ancestor with *mel* around 20 MYA (19), half the time separating *mel* from *vir* (Fig. 1b). Carriers of the *gt*^pse^ whole-locus ortholog exhibits reduced RV in both males and females, though to a lesser extent than carriers of *gt*^*vir*^ (Fig 2a, d). The reduction in RV by *gt*^*pse*^ is largely attributable to non-coding sequence (Fig 2f), and, like *gt*^*vir*^, there is also a coding contribution. Specifically, whereas the *gt*^*pse*^ coding shows reduced RV with its *gt*^*pse*^ noncoding region, the chimera carrying a *gt*^*mel*^ coding region does not (Fig 2a, c; Table S5). The set of experiments with *pse* and *vir* show striking parallels: strong contributions to reduced RV by the noncoding region; a contribution by the coding regions; and epistatic interaction between coding and noncoding for RV for *gt*^*vir*^ and possibly for *gt*^*pse*^ (Fig 2a,c).

### Functional divergence of closely related orthologs

*D. yakuba* (*yak*), *D. santomea* (*san*) and *D. erecta* (*ere*) belong to the same phylogenetic clade that has a common ancestor with *mel* 10 MYA (19) (Fig. 1b); one of them, *san*, produces viable hybrids with *mel*. Reduced RV is observed in two of the three species: *san* — *gt*^*san*^ whole locus (Fig 2d), *gt*^*san*^ eGFP-tagged whole locus, and *gt*^*san*^ coding-only (Fig. 2g, h); and *ere* — *gt*^*ere*^ eGFP-tagged whole locus. Thus, even on the relatively short timescale of 10MY separating this clade of species from *mel*, the experiments functionally distinguish their *gt* alleles from the *mel* ortholog.

### Species hybrids

Our viability assays thus far reveal functional differences between the *mel* allele and the *san, ere, pse* and *vir gt* orthologs. Unresolved is whether *yak*, the remaining species in our *gt* analysis, might also have functionally diverged from *mel gt*, albeit more subtly. We investigated this question with species hybrids. Crosses between *mel* females and *san* males produce sterile hybrid female progenies only. We have recently shown that *mel* Gt, differing by seven amino acid substitutions from *san* Gt (Table S4), causes reduced female viability in the hybrid (28). Acting on this finding, we tested additional *gt* transgene orthologs in *mel/san* hybrids by crossing *mel* females hemizygous for a transgene to *san* males (Fig. S4). In this cross, RV is estimated from the number of hybrid F1 flies carrying either the *gt* transgene or a control chromosome bearing no transgene. Chimeric transgenes whose coding regions have been replaced by *yak, ere* or *san* orthologs, under the regulatory control of *mel* noncoding region, all eliminate the deleterious effect of *mel* Gt in hybrid females (Fig 2i). In this sensitized hybrid genetic environment, we are thus able to place the functional divergence in the protein leading to reduced hybrid viability caused by *mel* Gt to changes on the phylogenetic branch leading to *mel* itself.

### Functional divergence of the *san gt* protein

Transgenic *yak* and *san gt* proteins under the regulatory control of *mel* noncoding sequence have significantly different RV in *mel* (Fig. 2h). This means that our experiments have detected *gt* functional divergence at every timescale separating the six species employed in our analysis. *san* and *yak* have a common ancestor estimated to be only 1.2 MYA (20), and Gt^*yak*^ and Gt^*san*^ proteins differ by only two substitutions — A351V and +4Q — (Fig. 3a). Gt^*yak*^ is identical at both sites to the allele in *D. teissieri*, the closest outgroup species, indicating that the Gt^*yak*^ carries both ancestral states (Fig. S3). We confirmed that the *yak* and *san* Gt alleles used in this experiment are both common alleles, not unique to specific populations of either species (Fig. S5, S6). With only two substitutions, both of which occurred in *san*, there are only two possible intermediate evolutionary paths. We investigated both of them in *mel* with eGFP-tagged transgenes carrying the two single-substitution genotypes under the regulatory control of *mel* noncoding sequence. Our RV assay reveals a significant increase in RV for the +4Q substitution alone and a significant decrease for the A351V substitution alone. Together, the two substitutions produce the most severe decrease in RV (Fig. 3b). Thus, both single substitutions have significant RV effects, the two possible trajectories differ significantly, and there is sign epistasis along one path. To summarize, no individual substitution in this sensitized experimental system is functionally inert, and together the two substitutions interact at the level of RV.

**Figure 3.**
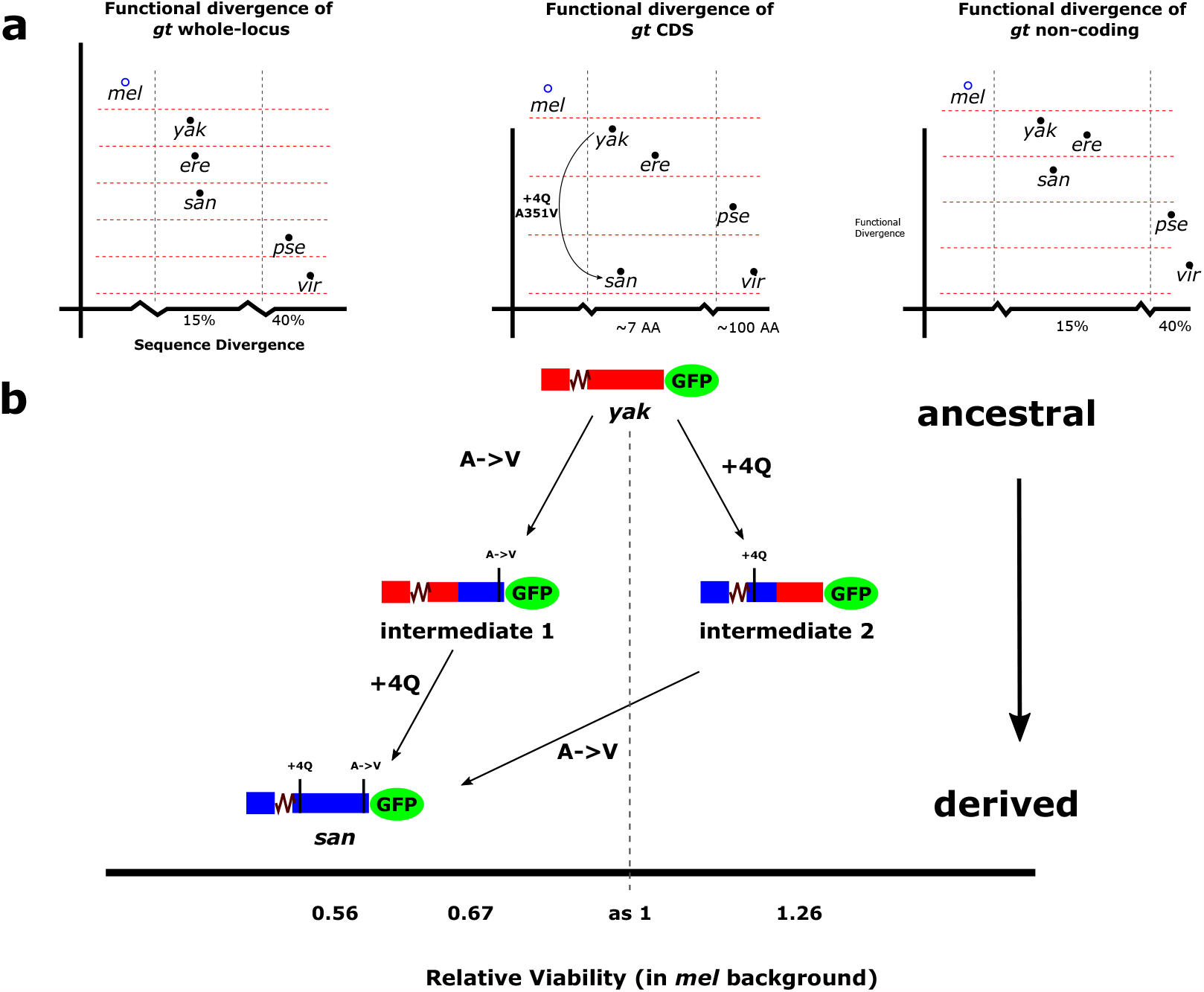
The two possible evolutionary intermediates of *gt* coding substitutions between *yak* and *san* have distinct (and opposite) relative viability effects in *mel*. **a**, Summary of *gt* functional divergence inferred from whole-locus or chimeric transgenes. Horizontal dashed lines demarcate distinguishable orthologs. **b**, Reconstructed ancestral states of *yak/san gt* protein and alternative evolutionary paths of divergence. Both alternative trajectories require non-neutral intermediates. Data in Table S6.

## Discussion

Our experimental results reveal a continuous process of functional divergence across *Drosophila* phylogeny and timescales, a sharp refutation of interchangeability of genes regulating evolutionarily conserved developmental processes. Our sensitized assays, employing appropriate transgene controls inserted into the same chromosomal docking site, identified functional divergence attributable to both coding and noncoding regions along nearly every branch of the phylogeny (Fig 4). Most published studies of orthologous gene function (see Table S1), i.e., interchangeability, investigate only protein coding regions and not the whole locus. Our study reveals limitations in this approach: none of protein coding regions alone from any of the five species significantly reduced RV when driven by the *mel* noncoding region (Fig. 2b, e). Our finding points to the whole locus as an integral target for *gt* functional evolution.

**Figure 4.**
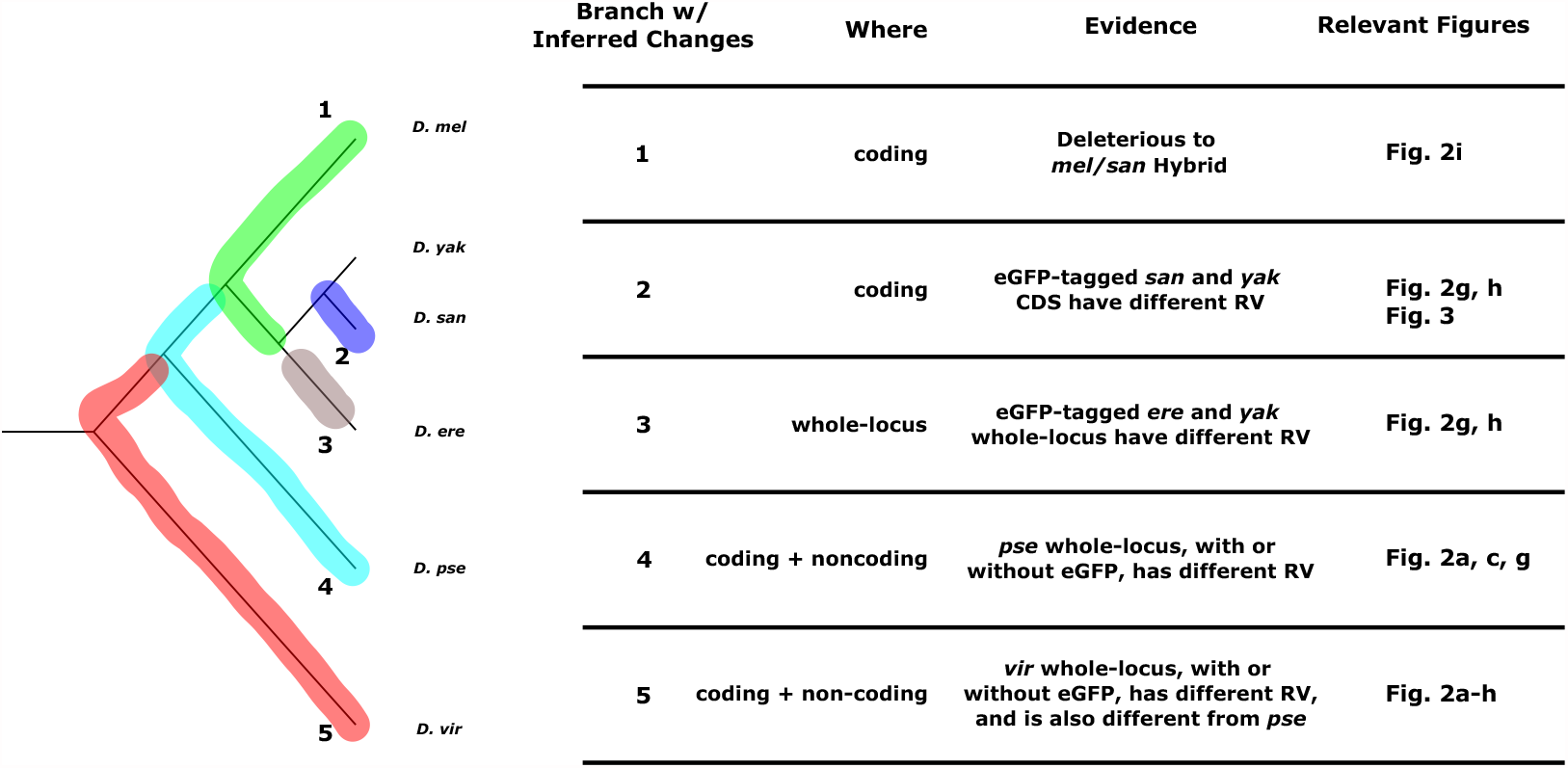
Continual functional divergence of the *gt* locus in *Drosophila*. A parsimonious reconstruction of functional changes in *gt* mapped onto phylogenic branches, marked by different colors. Experimental evidence for viability differences attributable whole-locus or chimeric transgene divergence is listed alongside each branch. Functional divergence in noncoding regions inferred from species-specific epistasis is not included because of uncertainty about branch assignment.

Functional changes in *gt* are relevant to understanding the genetics of interspecies hybrid incompatibility. Here, our experiments amplified on a recent finding that identified the protein coding region of *gt* in causing inviability in hybrids between *mel* and *san* (28). *We mapped those differences to substitutions in the phylogenetic branch leading to mel* from its common ancestor with *yak/san/ere*. This functional divergence is not unique to this single lineage, however, but rather is one instance of a continuous process of functional divergence across *Drosophila* phylogeny (Fig. 4). Hybrid incompatibility generally results from functional divergence of two interacting genes, one in each of two species, which when brought together in a hybrid, fail to function properly. The functional divergence in *gt*^*mel*^ protein is, therefore, likely accompanied by similar functional divergence in one or more interacting partners in *san*. In a search for a partner to *gt*, we discovered that orthologs of the gap gene *tailless* from *mel* and *san*, like *gt*, also differ in their effects on viability in the hybrid (28). In a broader context, continuous functional evolution of *gt*, as documented here, may be representative of other “conserved” genes in the gap gene network, and illustrative of the process of rapid molecular evolution leading to hybrid incompatibility.

We believe our unequivocal findings in experimental assays — a rapid, continuous process of *gt* functional evolution in coding and noncoding regions — are relevant to understanding population genetic mechanisms governing *gt* evolution. In general, one expects natural selection to be many orders of magnitude more sensitive to the fitness effects of subtle functional changes than those that can be measured in our laboratory experiment. In this context, no organism has received more attention than *Drosophila* in a quest to understand the extent of adaptive evolution driving gene and genome evolution, and there is now near-universal agreement that natural selection is the predominant driving force in these large-population-size species (29, 30). Our findings suggest that the very same mechanism — natural selection — may be responsible for the continuous pace of *gt* functional evolution. Especially illuminating are the two coding substitutions between the very closely related species *yak* and *san*: both evolutionary intermediates, and the combination of substitutions together, all exhibit significant viability effects in a *mel* genetic background. The fact that these *gt* intermediates are distinctly different in the same genetic background suggests to us that natural selection is likely to have been involved in their substitution, even if the fitness effects are smaller in their ancestral backgrounds.

Under this mechanistic framework, the conservation of gap gene network output is achieved both by selective constraint acting on the network, as well as by a continuous process of functional refinement to individual genes and their cross-regulatory interactions. The continuous functional divergence of gap genes also gives rise, inevitably, to changes in the detailed molecular mechanisms by which the network directs pattern formation, a characteristic of developmental system drift. Our discovery of rapid functional divergence of *gt* requires reassessment of the tempo and mode of molecular evolution of regulatory genes belonging to conserved developmental systems.

## ACKNOWLEDGEMENTS

This work was supported by the National Science Foundation under award 1916895 to M.K., and by NIGMS R01 GM125715 to D.R.M. We thank John Reinitz, David Stern, Urs Schmidt-Ott, Manyuan Long and Andreas Wagner for comments, and the University of Chicago DNA sequencing & Genotyping Facility for Sanger sequencing.

## AUTHOR CONTRIBUTIONS

W.C. and M.K. conceived the project and shared in writing the manuscript. W.C. performed the transgenic and viability experiments and carried out the analysis. D.R.M. shared in discussions about the project, provided *D. santomea* stocks, gave advice about conducting the interspecific hybrid experiment, and contributed to editing the manuscript.

